# The role of the meningeal lymphatic system in local inflammation and trigeminal nociception implicated in migraine pain

**DOI:** 10.1101/2020.04.25.060939

**Authors:** Nikita Mikhailov, Kseniia Koroleva, Ali Abdollahzadeh, Raisa Giniatullina, Oleg Gafurov, Tarja Malm, Alejandra Sierra, Jussi Tohka, Francesco Noe, Rashid Giniatullin

## Abstract

**Background:** A system of lymphatic vessels has been recently characterized in the meninges, with a postulated role in ‘cleaning’ the brain via cerebral fluid drainage. As meninges are the origin site of migraine pain, we hypothesized that malfunctioning of the lymphatic system should affect the local trigeminal nociception. To test this hypothesis, we studied nociceptive and inflammatory mechanisms in the meninges of K14-VEGFR3-Ig mice lacking the meningeal lymphatic system.

**Methods:** We recorded the spiking activity of meningeal afferents and estimated the local mast cells infiltration, calcitonin gene-related peptide (CGRP) and cytokine levels (basal and stimulated), as well as the dural trigeminal innervation in freshly-isolated hemiskull preparations from K14-VEGFR3-Ig (K14) or wild type C57BL/6 mice (WT).

**Results:** We found that the meningeal level of CGRP and of the pro-inflammatory cytokines IL12-p70 and TNFα (implicated in migraine) were reduced in the meninges of K14 mice. On the contrary, in the meninges of K14 mice, we found an increased level of the mast cell activator MCP-1 and, consistently, a larger number of dural mast cells. The other migraine-related pro-inflammatory cytokines did not differ between the two genotypes. The patterns of trigeminal innervation in meninges remained unchanged and we did not observe alterations in basal or ATP-induced nociceptive firing in the meningeal afferents.

**Conclusions:** In summary, the lack of meningeal lymphatic system does not induce migraine-like nociceptive state *per se*, but leads to a new balance between pro- and antiinflammatory factors implicated in migraine mechanisms.

## 1 Introduction

Migraine is a complex neurological disorder, which originates from the meninges (Bolay et al., 2002; Olesen et al., 2009). Meninges are densely innervated by somatic trigeminal (Ramachandran, 2018; Zakharov et al., 2015) and parasympathetic innervation (Delépine and Aubineau, 1997), which runs along and in contact with meningeal blood vessels (Ebersberger et al., 2006). It has been proposed that so-called sterile neurogenic inflammation (Moskowitz, 1993, 1984; Moskowitz et al., 1979) in the meninges is an important initiator of the pathophysiology of migraine. The initial step in neurogenic inflammation is the activation of the trigeminal nociceptive system in meninges (Moskowitz, 1993). Indeed, trigeminal neurons release the calcitonin gene-related neuro-peptide (CGRP), which activates dural mast cells (Ottosson and Edvinsson, 1997), promotes sensitization of trigeminal afferents (Cady et al., 2011; Giniatullin et al., 2008), and induces local vasodilation (Goadsby et al., 1988). Vasodilation, in turn, activates mechanosensitive receptors, expressed in trigeminal nerves within the meninges (Mikhailov et al., 2019). Trigeminal neurons can also be activated by mast cells-secreted serotonin (Kilinc et al., 2017), and by extracellular ATP, which is released from endothelial and mast cells, as well as from neurons, upon stimulation of the migraine mediator CGRP (Koroleva et al., 2019; Suleimanova et al., 2020; Yegutkin et al., 2016). Finally, the release of acetylcholine (ACh) by parasympathetic fibers results in meningeal mast cell degranulation (Shelukhina et al., 2017), which activates the trigeminal nociceptive system. In addition, endothelial (Loberg et al., 2006) and mast cells (Mukai et al., 2018) release various pro-inflammatory cytokines as potential contributors to local inflammation.

The presence of these diverse endogenous agents in the meninges suggests the existence of regulatory mechanisms, which modulates the local levels of pro-inflammatory mediators and minimize the activity of nociceptive fibers. The recently characterized meningeal lymphatic system (Antila et al., 2017; Aspelund et al., 2015; Louveau et al., 2015) **(Figure 1)** has been suggested to play a role in the clearance of the central nervous system (CNS) from toxic molecules, and to be involved in the pathogenesis of different neurodegenerative diseases (Kwon et al., 2019; Noé and Marchi, 2019; Patel et al., 2019; Rasmussen et al., 2018; Tamura et al., 2019). We hypothesized that the meningeal lymphatic vessels (mLVs) can play a similar role also in the meninges, clearing the pro-inflammatory mediators. We also hypothesize that dysfunction in the lymphatic drainage can trigger migraine.

**Figure 1.**
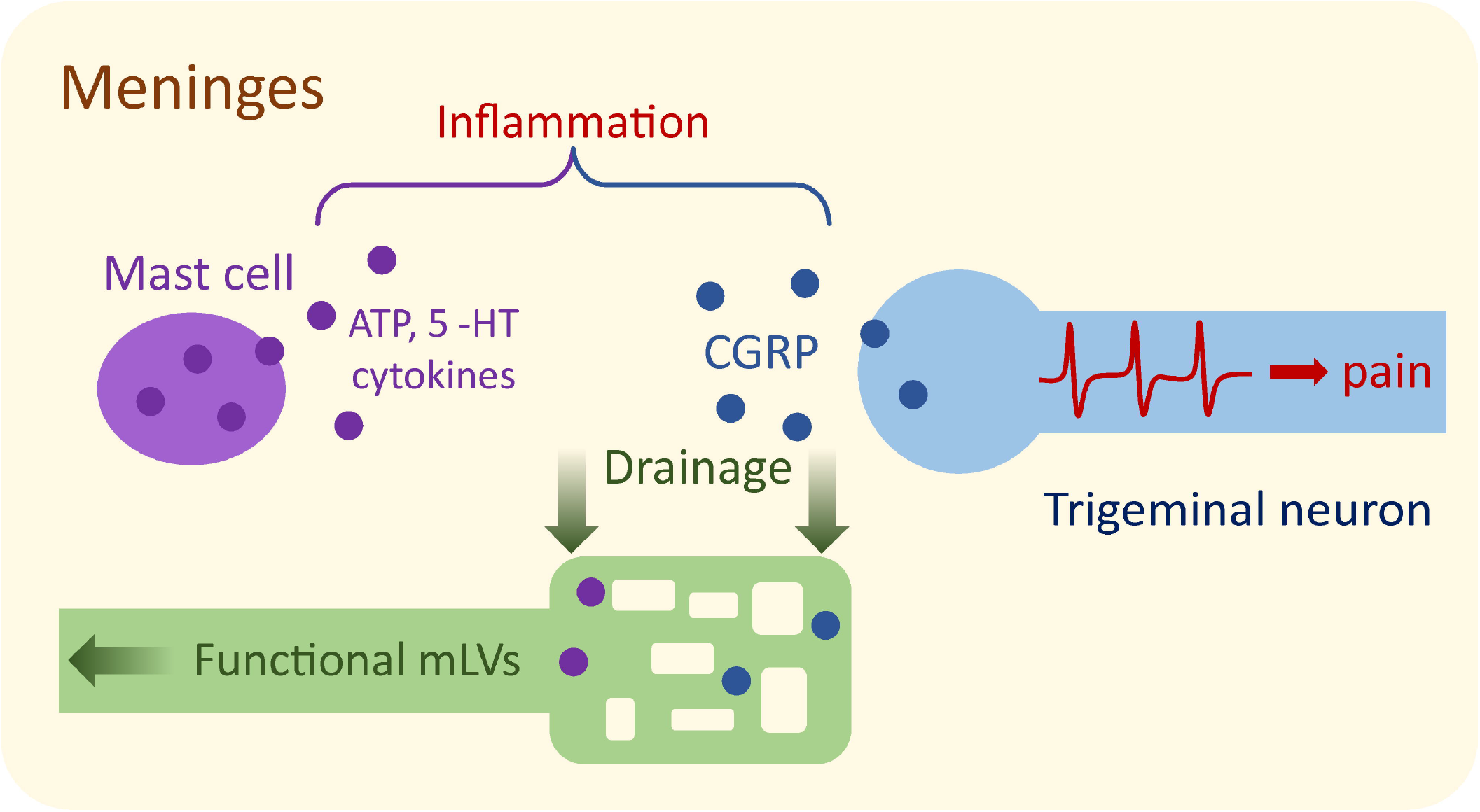
Migraine-related cell types and active mediators present in meninges. The main migraine mediator CGRP is released from the meningeal trigeminal nerve fibers, whereas ATP, serotonin (5-HT) and cytokines are released from local immune cells, including mast cells. We propose that the local lymphatic system, which supposedly contributes to meningeal drainage, maintain local homeostasis preventing the accumulation of pro-inflammatory and pronociceptive compounds.

## 2 Methods and materials

### 2.1 Animals

We used adult (30 – 40 g) male and female transgenic K14-VEGFR3-Ig (K14) mice lacking the meningeal lymphatic system (Aspelund et al., 2015; Mäkinen et al., 2001) and wild-type C57BL/6J littermates mice (WT). Total number of animal used for this study was 38 WT and 37 TG animals. Animals were housed at the Lab Animal Center of the University of Eastern Finland under following housing conditions: 12 hours dark/light cycle, grouped housing, given ad libitum access to food and water, 22°C ambient temperature. All procedures with animals were conducted in accordance with the European Community Council Directive 2010/63/EU. Experimental protocols involving the usage of animals were approved by the Animal Care and Use Committee of the University of Eastern Finland (license EKS-004-2014) and by the National Animal Experiment Board (ESAVI-2017-008787).

### 2.2 Hemiskull preparation for ex vivo experiments

For the *in vitro* and *ex vivo* experiments (mast cell staining, CGRP release assay, cytokine release assay, and electrophysiology), we used hemiskull preparations with preserved meninges and meningeal trigeminal innervation (Shatillo et al., 2013; Zakharov et al., 2015). For electrophysiology and CGRP release assay, animals we sacrificed with CO2 without anaesthesia. For cytokine release assay and mast cell staining, animals were anesthetized with intraperitoneal injection of 250 mg/kg tribromoethanol (Avertin, Sigma-Aldrich, USA) and then underwent transcardial perfusion: for mast cell staining, we perfused with normal saline (6 minutes at 6 ml/min), followed by 4% paraformaldehyde (PFA) (20 minutes at 3 ml/min); for cytokine release assay, animals were just perfused with normal saline (4 min at 20 ml/min). In order to prepare the hemiskulls for subsequent analyses, after decapitation, skin, muscles, and connective tissues, as well as brain were removed with particular care to not touch and damage the meninges. Thereafter, hemiskulls were isolated by cutting the skull along the sagittal axis (*i.e.*, two hemiskulls were prepared from each animal).

### 2.3 Mast cell staining

Hemiskulls of WT and K14 animals were prepared as described above, washed three times for 20 minutes in PBS (phosphate buffered saline), and stained for 60 minutes with 0.1 % toluidine blue. Stained meninges were immediately imaged, using a Zeiss Axio Zoom.V16 stereomicroscope (Carl Zeiss AG, Germany), at 30x magnification. We imaged two identical regions-of-interest (ROIs) from each hemiskull: one including the superior sagittal sinus (SSS; at the border of the image) and the adjoining one, partially overlapping (approximately 10% of area) with the previous ROI. Size of each ROI was 3.7×2.4 mm^2^. In total, four images were acquired from each mouse. From each ROI, the number of stained mast cells was calculated using ImageJ 1.51v software (National Institute of Health, USA).

### 2.4 CGRP release assay

The CGRP release assay was conducted on hemiskulls of WT and K14 animals as previously described (Mikhailov et al., 2019). We used a CGRP enzyme immunoassay kit (EIA kit, SPIbio, France) to measure CGRP release from hemiskull preparations. The levels of CGRP were assessed under control and KCl-treated conditions.

First, fresh hemiskull preparations from WT and K14 animals were perfused for 40 min with artificial cerebrospinal fluid (aCSF in mM: 119 NaCl, 30 NaHCO3, 11 glucose, 2.5 KCl, 1 MgCl2, 1 NaH2PO4, 2 CaCl2, adjusted pH level = 7.4) at room temperature (RT). Then, hemiskulls were washed four times with 150 μl aCSF for 15 minutes at 37°C to stabilize the basal condition. Samples of aCSF (100 μl each) from the last two washes were collected to assess the basal level of CGRP (control conditions). After the last wash, hemiskulls were bathed with an aCSF solution containing 30 mM KCl for 15 minutes. Thereafter, a 100 μl sample was collected to assess CGRP under KCl-treated condition.

Samples were freezed with liquid nitrogen in tubes containing EIA buffer with peptidase inhibitors. Assay protocol was carried out in accordance with instructions of manufacturer. In brief, 100 μl of CGRP standard (for calibration curve) or sample were mixed with 100 μl of anti-CGRP AChE tracer in a well plate, preliminary washed 5 times with washing buffer. After incubating the plates at 4 °C for 16-20 h, supernatant was removed, and plates were washed 6 times with washing buffer. Finally, 200 μl of Ellman’s reagent was added. Optical density was measured at 405 nm using microplate reader (Wallac VICTOR2™, PerkinElmer, USA).

### 2.5 Cytokine release assay

For cytokine release assay, we used hemiskull preparations from both WT and K14 mice. One hemiskull/animal was used to assess the release of cytokine under the control condition, while the contralateral was used to measure the release of cytokine under treatment condition (100 μM benzoyl ATP, BzATP). After the collection of hemiskulls, all procedures were performed at 37°C. Hemiskulls were bathed in 150 μl of either aCSF or aCSF + BzATP for three and a half hours. aCSF samples (50 μl) were taken to assess the level of cytokines in control (treated with aCSF only) or stimulated (aCSF + BzATP) conditions. The more stable ATP-analogue BzATP was used for the cytokine release assay, due to the long treatment time (3½ h) and because of fast degradation of ATP (Yegutkin et al., 2016). The levels of aCSF in the hemiskulls were monitored throughout the incubation period. If needed, we refilled the hemiskull preparation with the respective treatment solution to compensate for evaporation. To assess the level of cytokines (IL-6, IL-10, MCP-1, TNFα, IFNγ, IL-12p70), we used the CBA Mouse Inflammation Kit (Cytometric Bead Array, BD Biosciences, New Jersey, USA). All procedures (sample and standard beads preparation) were performed following instructions provided by the manufacturer. Data were acquired on the CytoFLEX S cytometer (Beckman Coulter Inc., USA) and analyzed with the Flow Cytometric Analysis Program (FCAP) Array software (BD Biosciences, USA).

Due to the low concentration of IL-12p70 and IFNγ, in some samples it was not possible to quantify the exact concentration with the abovementioned method. Thus, for these cytokines final *n* was lower than the total number of animals used for the assay.

### 2.6 Electrophysiology

Direct recording of action potentials from the trigeminal nerve afferents innervating meninges was conducted as described earlier (Shatillo et al., 2013; Zakharov et al., 2015). Briefly, hemiskulls of WT and K14 animals were placed in a perfusion chamber with a constant flow (6 ml/min) of oxygenated aCSF. In order to access the nervus spinosus of the trigeminal nerve, innervating meninges, we made a small incision in the dura mater. Next, the nervus spinosus was sucked into a glass recording electrode filled with aCSF. We recorded 10 min of spontaneous activity to get a baseline and to assess the difference in basal activity between WT and K14 animals. Then, we applied 100 μM ATP for 10 min to stimulate nociceptive firing in the trigeminal nerve (Zakharov et al., 2016), followed by 10 min of washout. The recording was conducted with a digital amplifier (ISO 80, WPI Inc., USA) with bandpass 300 Hz – 3 kHz, gain 10000. Signals were digitized at 125 kHz using a NIPCI 6221 data acquisition board (National Instruments, USA) and visualized with WinEDR software (Strathclyde University, UK).

### 2.7 Meningeal nerve staining for β-tubulin III

For the meningeal nerve staining, meninges from PFA-fixed hemiskulls were isolated as previously described (Louveau et al., 2015), and stained. Briefly, meningeal whole-mount preparations were incubated with 1x phosphate-buffered saline (PBS) containing 2% normal goat serum (NGS), 1% bovine serum albumin (BSA), 0.1% Triton X-100, and 0.05% Tween 20 for 1 hour at RT. After that, meninges were incubated with rabbit anti-β-tubulin III (1:1000, Cat# T2200, Sigma-Aldrich, USA) overnight at 4°C in PBS containing 1% BSA and 0.5% Triton X-100. Whole-mounts were washed in PBS at RT (3x) followed by incubation with Alexa Fluor 488-conjugated goat anti-rabbit IgG antibody (1:500, Cat# A11008, Invitrogen, USA) for 1 hour at RT in PBS with 1% BSA and 0.5% Triton X-100. After further washing in PBS (3x) and in phosphate buffer (PB; 2x), meninges were mounted with VECTASHIELD mounting medium (Vector Laboratories, USA), including DAPI, and coverslipped.

Images were acquired using a Leica TCS SP8 X confocal system (Leica Microsystems, Germany) using the LAS X software. Images of the region of interest, adjacent to the middle meningeal artery, were acquired with a 20x objective with 0.75 numerical aperture (NA), with an in-plane pixel size of 1.14×1.14 μm^2^ and a z-step of 1 μm. The full images were created by merging 4×4 tile scans, covering a total area of 2034×2034 μm^2^.

### 2.8 Axon segmentation and innervation analysis

We developed a semi-automated segmentation technique to annotate axons and analyze the innervation in acquired 3D confocal microscopy images. An experienced researcher (AA) inserts two points along the desired axon on the 2D maximum projection of a 3D confocal image to define the axonal centerline. Given the two points, a minimal path algorithm (Benmansour and Cohen, 2011) detects the centerline of that axon. We used active contours (Chan and Vese, 2001) to segment the desired axon, initiating from the detected axonal centerline. For each axon, we measured the axonal diameter and the length of the centerlines in 3D (Abdollahzadeh et al., 2019a). We also quantified an innervation complexity value for each meningeal tissue by forming a graph from the segmented axons (Abdollahzadeh et al., 2019b) and reported the number of axonal endpoints, as shown in Fig. 7a. We described the details of axon segmentation and the morphology analysis in Supplementary Materials (“axon segmentation and innervation analysis” section).

### 2.9 Statistical analysis

We performed statistical analysis using the GraphPad Prism 8.4.2 software (GraphPad Software, California, USA). The data are presented as mean ± the standard error of the mean (for two-way ANOVA) or as median with interquartile range (for Mann-Whitney U test and linear mixed effect model). We set the alpha-threshold defining the statistical significance as 0.05 for all analyses. Mast cell abundance as well as axon segmentation and innervation analyses data were analysed using Mann-Whitney U test. CGRP release assay data was analysed with repeated measures two-way ANOVA. Cytokine release assay data were analysed by Mann-Whitney U test or by matched two-way ANOVA. A linear mixed effect model was used to evaluate the differences in the electrophysiology data. Bonferroni correction was used to adjust p-values in multiple comparison. Please see figure legends for the details of the used test in each analysis.

## 3 Results

### 3.1 Elevated number of mast cells in meninges of mice without lymphatic system

Meningeal mast cells recently emerged as important players in the initiation of a migraine attack (Giniatullin et al., 2019; Kilinc et al., 2017; Koyuncu Irmak et al., 2019; Levy, 2009). As the mLVs are important for the movement of immune cells (Ahn et al., 2019; Louveau et al., 2015), we expected that the lack of mLVs could affect the local accumulation of mast cells. We used the toluidine blue labelling to characterize the profile of meningeal mast cells.

In WT animals a substantial portion of mast cells was located along the meningeal blood vessels (**Figure 2A**), where the mLVs are located (Antila et al., 2017; Aspelund et al., 2015). A similar pattern of mast cell positioning was observed in K14 animals (**Figure 2B**). K14 animals showed a non-significant trend towards increased number of meningeal mast cells (**Figure 2.C**). Since an increase in the number of mast cells may lead to a more abundant release of local mediators (Giniatullin et al., 2019; Levy, 2009; Suleimanova et al., 2020), resulting in meningeal inflammation and in the stimulation of the trigeminal afferents (Kilinc et al., 2017; Koroleva et al., 2019), we analyzed the levels of the different migraine-associated pro-inflammatory molecules and the activation of the trigeminal nerves in the K14 and their WT littermates mice.

**Figure 2.**
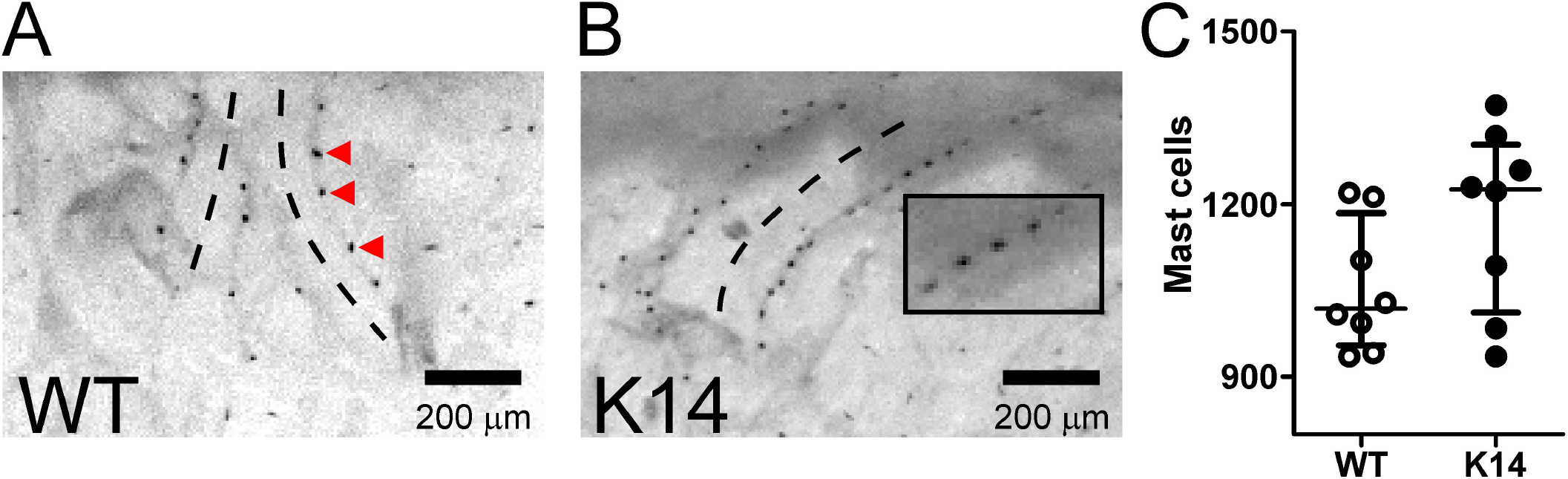
Dural mast cells in WT and K14 mice lacking the lymphatics. Mast cells localize in meninges in both WT **(A)** and K14 mice **(B)**. Note the presence of mast cells (red triangles) aligned along the blood vessels (outlined with black dashed curves). **(C)**Mast cells were counted in WT and K14 animals. Transgenic animals show a non-significant trend towards a larger number of mast cells (1055 ± 40 cells in WT *vs.* 1177 ± 56 cells in K14, n = 8/experimental group, Mann-Whitney U = 16.5, p = 0.1089 by Mann-Whitney U test). Scale bar = 200 μm.

### 3.2 Altered cytokine release in meninges

We first measured the level of cytokines (IL-6, IL-10, IL-12p70, MCP-1, TNFα, and IFNγ) released into meninges in basal conditions (**Figure 3**). The level of the pro-inflammatory MCP-1 was significantly higher in K14 animals compared to their WT controls (**Figure 3C**). On the contrary, the basal level of IL-12p70 (also a pro-inflammatory cytokine) was significantly decreased in K14 animals compared to WT littermates (**Figure 3E**). No differences in the levels of all the other measured cytokines were observed in basal conditions between the two genotypes (**Figure 3**).

**Figure 3.**
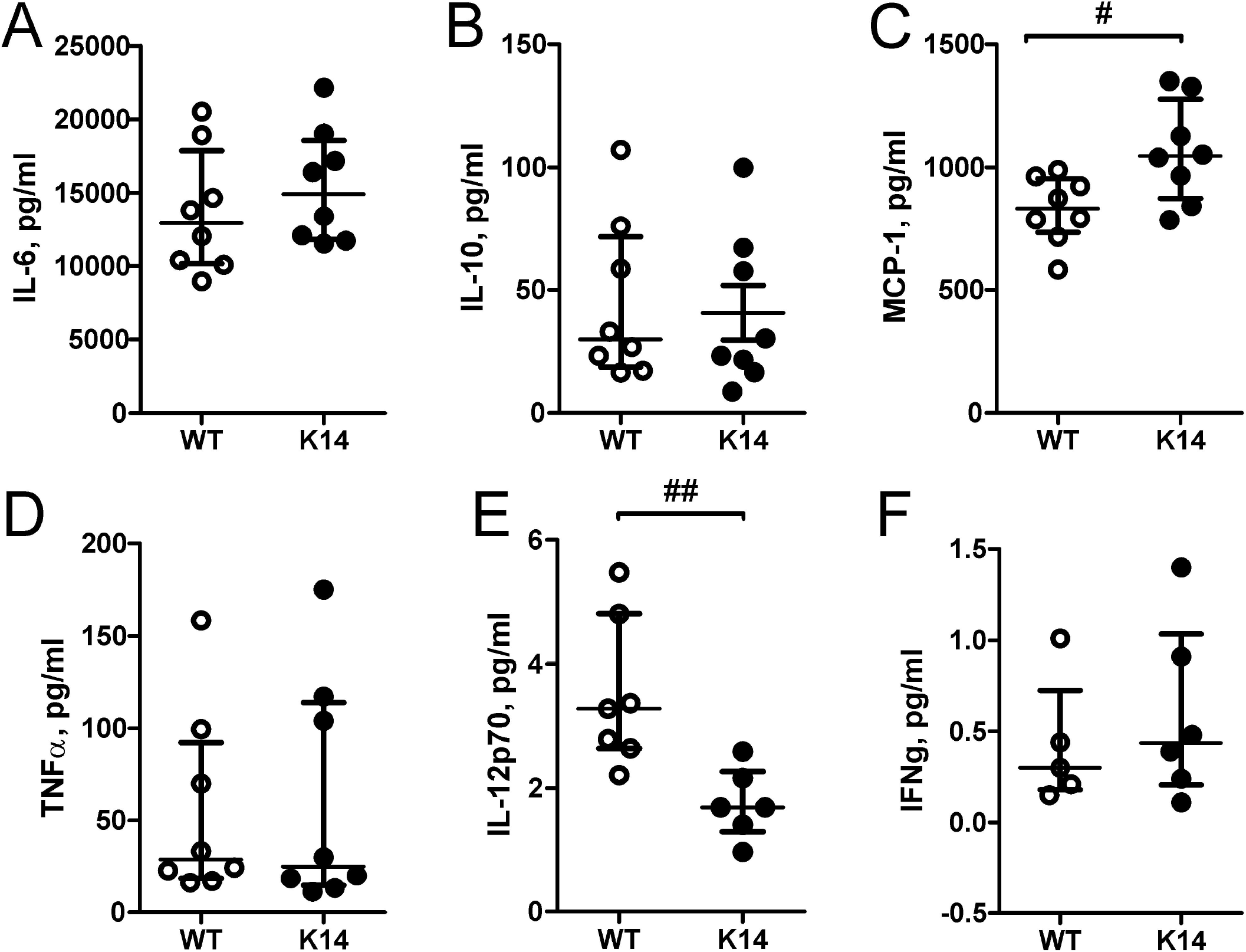
Baseline cytokines release from meninges. Concentrations (pg/ml) of following cytokines are depicted: IL-6 **(A)**, IL-10 **(B)**, MCP-1 **(C)**, TNFα **(D)**, IL-12p70 **(E)** and IFNγ **(F)**. In K14 animals, we found a significant increase in the levels of MCP-1 **(C)**(829 ± 48 pg/ml in WT *vs.* 1061 ± 72 pg/ml in K14, Manh-Whithey U = 11, #p = 0.0281, Mann-Whitney U test), whereas the levels of IL-12p70 **(E)** were significantly lower compared to the one measured in WT littermates (3.5 ± 0.5 pg/ml in WT *vs.* 1.8 ± 0.2 pg/ml in K14, Mann-Whitney U = 1.0, ##p = 0.0023, Mann-Whitney U test). No difference was found for other interleukines. Number of replicates for IL-6, IL-10, MCP-1 and TNFα analysis n = 8/experimental group; for IL-12p70 analysis: n = 7 and 6 for WT and K14, respectively; for IFNγ analysis: n = 5 and 6 for WT and K14, respectively.

To mimic inflammatory conditions in the meanings, we used BzATP, a stable ATP analogue (Baraldi et al., 2005), to stimulate the pro-inflammatory P2X7 receptors (Takenaka et al., 2016) in the dural immune cells, including mast cells (Nurkhametova et al., 2019). Then, we measured the cytokine release from meninges of both WT and K14 animals. Stimulation with 100 μM BzATP significantly increased the levels of IL-10 and TNFα (**Figure 4B, D)**. Interestingly, while the IL-10 increase did not differ between the WT and K14 mice, BzATP-stimulated release of the pro-inflammatory cytokine TNFα was significant exclusively in the WT genotype (**Figure 4D**). BzATP stimulation did not alter the levels of other measured cytokines.

**Figure 4.**
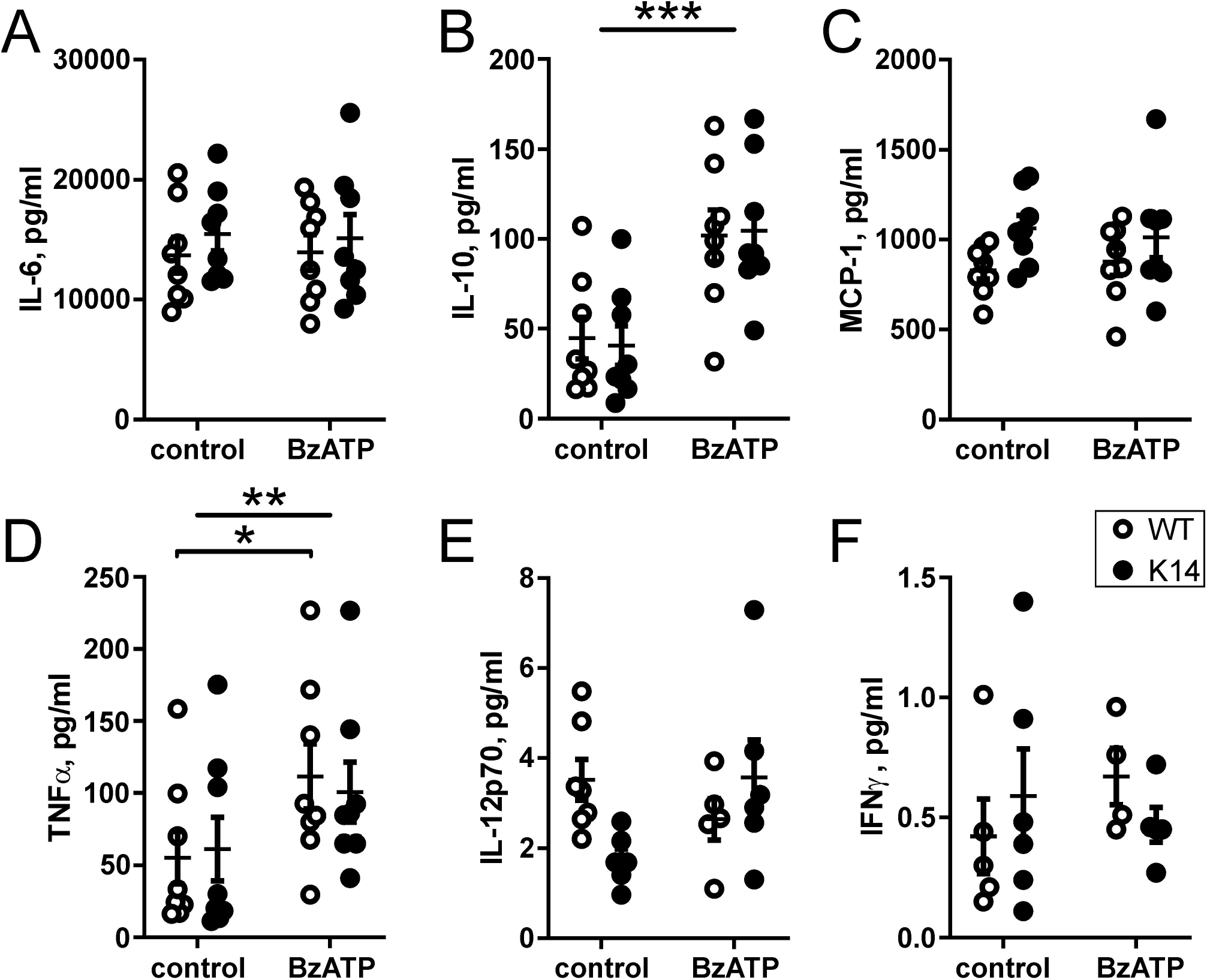
Cytokines release from meninges after BzATP stimulation. Scatter plots reporting the concentrations of cytokines released from meningeal preparation after stimulation with BzATP: IL-6 **(A)**, IL-10 **(B)**, MCP-1 **(C)**, TNFα **(D)**, IL-12p70 **(E)** and IFNγ **(F)**. BzATP induced an increase in the release of IL-10 **(B)**(treatment effect WT-K14-control *vs.* WT-K14-BzATP, F(1, 14) = 35.75, ***p < 0.0001 by matched two-way ANOVA) and TNFα **(D)**(WT-K14-control *vs.* WT-K14-BzATP, F(1, 14) = 12.43, **p = 0.0034 by matched two-way ANOVA). Post-hoc comparisons revealed that IL-10 increase did not differ between WT and K14, whereas BzATP-stimulated release of TNFα was significant only in the WT mice (55 ± 18 pg/ml in WT-control *vs.* 112 ± 23 pg/ml in WT-BzATP, t(14) = 2.933, *p = 0.0218 by Bonferroni post-hoc test). Number of replicates for IL-6, IL-10, MCP-1, TNFα analysis: n =8/experimental group; for IL-12p70 analysis: n = 7 and 6 (control) and n = 5 and 6 (BzATP) for WT and K14, respectively; for IFNγ analysis: n = 5 and 6 (control) and n = 4 and 5 (BzATP) for WT and K14, respectively.

### 3.3 Decreased CGRP release in meninges of mice lacking lymphatic system

Next, we measured the level of CGRP released in meninges in both basal and stimulated conditions (**Figure 5**). Basal CGRP levels, as measured in two consecutive samples (baselines, BL), were similar in both genotypes. Interestingly, stimulation with 30 mM KCl, which leads to CGRP release through the stimulation of peptidergic nerve fibers, induced a significantly lower increase of CGRP levels in K14 compared to WT hemiskulls (**Figure 5A**).

**Figure 5.**
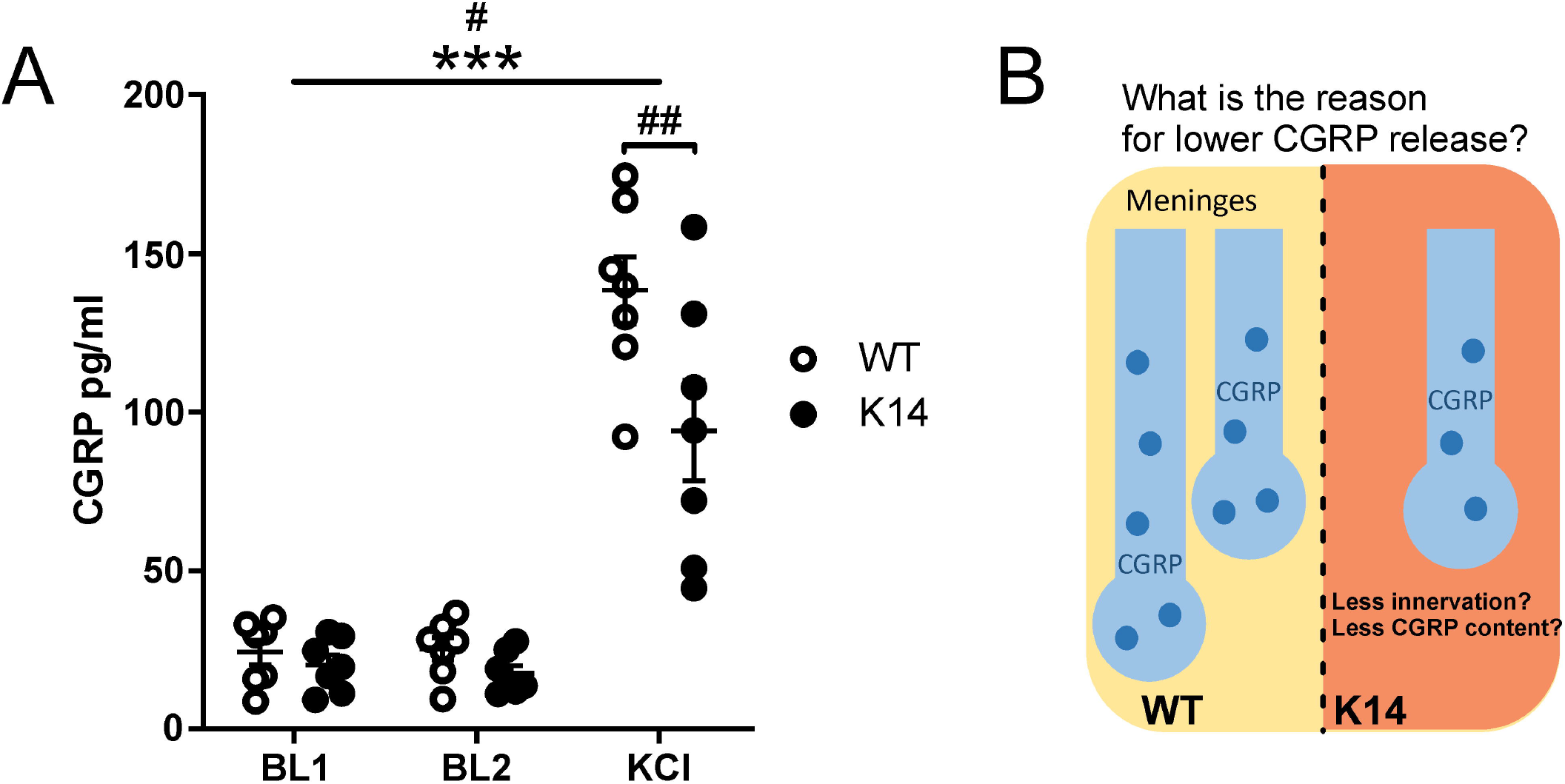
CGRP released from trigeminal nerve endings in the meninges, from hemiskull preparation. (A) CGRP release was not altered in K14 hemiskulls under basal conditions: 24 ± 4 pg/ml in WT-BL1 *vs.* 20 ± 3 pg/ml in K14-BL1, and 25 ± 3 pg/ml in WT-BL2 *vs.* 17 ± 2 pg/ml in K14-BL2 (n = 7/experimental group, p > 0.05 by matched two-way ANOVA with Bonferroni post-hoc test). The application of 30 mM KCl triggered a significant increase in the CGRP level in both genotypes (treatment effect: F(2, 24) = 103.3, ***p < 0.0001 by matched two-way ANOVA, n = 7/experimental group). Overall CGRP release was higher in WT animals (genotype effect: F(1, 12) = 6.025, #p = 0.0303 by matched two-way ANOVA). Post-hoc comparisons revealed that the concentration of the released CGRP under the stimulation was significantly lower in the hemiskulls lacking the lymphatic system compared to the WT preparations (94 ± 16 pg/ml in K14-KCl *vs.* 138 ± 11 pg/ml in WT-KCl; t(36) = 3.816, ##p = 0.0015 by Bonferroni post-hoc test, n = 7/experimental group). **(B)** We propose two possible explanations for the altered CGRP release: (1) lower density of trigeminal afferents releasing CGRP, or (2) lower expression of CGRP.

### 3.4 Spontaneous and ATP-induced spiking activity in meningeal nerves of mice lacking meningeal lymphatic system

To address the state of trigeminal nociception in animals without the lymphatic system, we recorded spiking activity from meningeal afferents. First, we recorded spiking activity in trigeminal nerve fibers in hemiskulls under basal conditions. Thereafter, we stimulated nerve terminals with ATP, a powerful trigger of local nociception (Koroleva et al., 2019; Yegutkin et al., 2016) (**Figure 6A**). In both genotypes, we observed a comparable increase in the nociceptive-related firing after ATP stimulation (**Figure 6B**).

**Figure 6.**
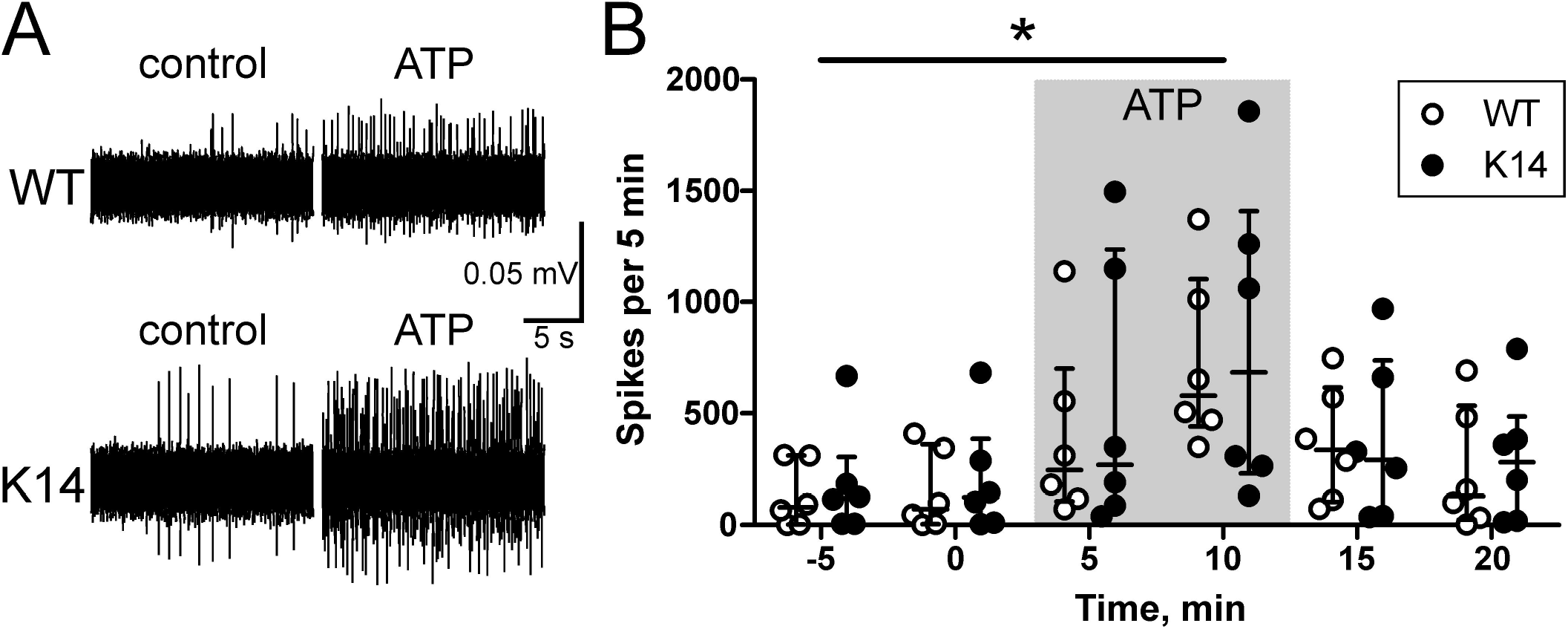
Direct spikes were recorded in WT and K14 hemiskulls under basal conditions and ATP stimulation. (A) Sample traces of nociceptive recordings in WT and K14 animals under basal and stimulated conditions. **(B)**The plot represents number of nociception-related spikes recorded in 5 minutes (median with interquartile range) under basal conditions, during the application of 100 μM ATP, and during the following washout. ATP induced a significant increase in nociception activity in both genotypes (baseline *vs.* ATP, t(50) = −2.65, * p = 0.0107 by linear mixed-effects model; n = 6/experimental group).

### 3.5 Unchanged patterns of meningeal innervation

To evaluate the density of trigeminal innervation in hemiskull preparations from WT and K14 animals we stained meningeal preparations with the antibodies specific to the neuronal marker β-tubulin III (**Figure 7**) (Cáceres et al., 1986). After that, we applied a semi-automated segmentation method to evaluate the following parameters: total innervation length (indirect evaluation of the innervation density in the meninges) and number of terminal points (*i.e.*, trigeminal fiber terminal endings, likely releasing CGRP). None of the analyzed parameters showed a difference between the genotypes.

**Figure 7.**
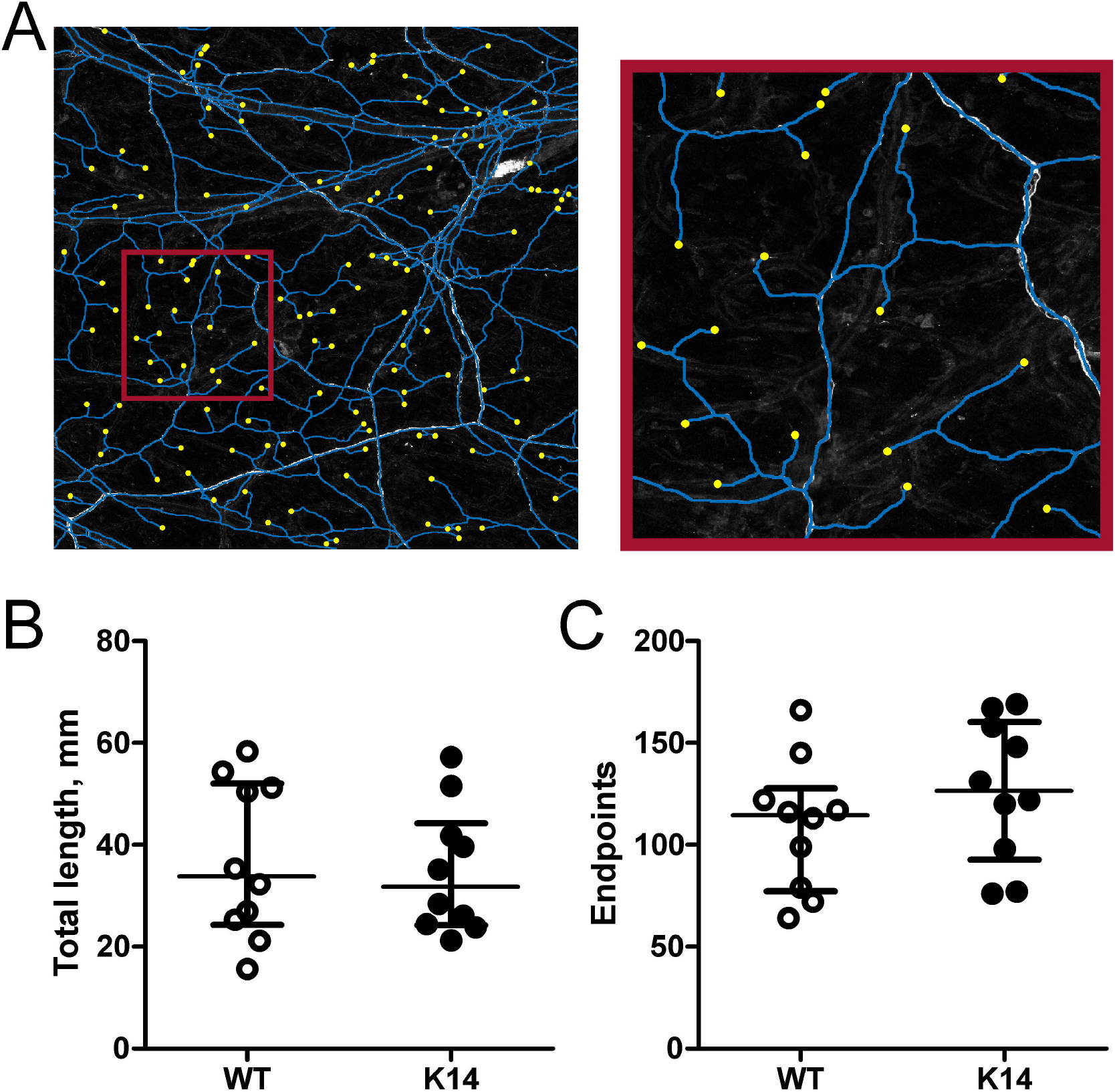
Segmentation analysis of meningeal innervation. (A) A representative image of meninges stained for β-tubulin III (white), segmented for morphological analyses. Blue lines represent axons, and yellow filled-circle represent terminal points (*i.e.*, CGRP-releasing trigeminal fiber terminal endings). **(B)**Scatter plot depicting the total axonal length (axons highlighted with blue on panel **A**): 37 ± 5 mm in WT *vs.* 35 ± 4 mm in K14, Mann-Whitney U = 47, p = 0.8534 by Mann-Whitney U test test. **(C)**Scatter plot depicting the number of terminal points (yellow dots on panel **A**): 109 ± 10 in WT *vs.* 127 ± 11 in K14, Mann-Whitney U = 32.5, p = 0.1972 by Mann-Whitney U test. We analyzed meningeal innervation from the right hemiskulls of 10 independent animals in each experimental group (WT and K14; one image per animal).

## 4 Discussion

Our study evaluated the role of the meningeal lymphatic system in the regulation of the neurochemical profile and functional properties of the meningeal trigeminovascular system (TVS), implicated in migraine pain. In K14 mice, lacking mLVs, we found that the stimulated release of CGRP, as well as the release of pro-inflammatory IL-12p70 and TNFα were decreased. On the contrary, we observed an increase in the levels of the pro-inflammatory cytokine MCP-1. The density of trigeminal meningeal innervation and nociceptive activity of nerve terminals were not altered in animals lacking meningeal lymphatic system. The main findings are summarized in **Figure 8**.

**Figure 8.**
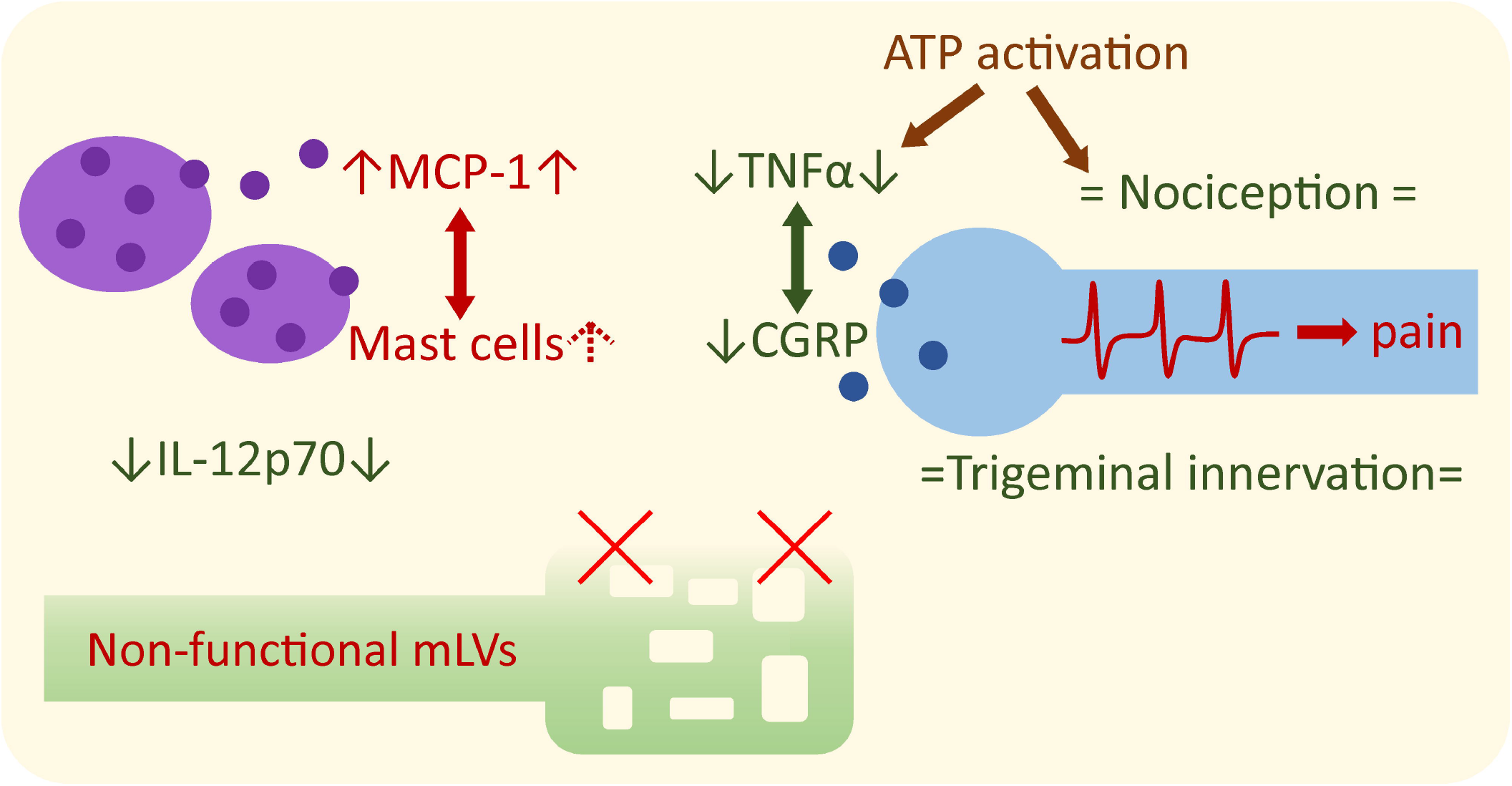
Experimental findings summary. Dysfunctional lymphatic system triggered alterations in migraine mediator levels in the meninges. Number of mast cells was increased, supported by increased level of mast cell-activating cytokine MCP-1. CGRP release was decreased, supported by decreased level of TNFα, a cytokine triggering CGRP transcription. The level of the pro-inflammatory IL-12p70 was decreased as well. Trigeminal innervation of the meninges and trigeminal nociception were not changed. Red indicates pro-inflammatory and pro-nociceptive changes, green indicates anti-inflammatory and anti-nociceptive changes. Arrow up (↑) indicates an increase in the parameter, arrow down (↓) – decrease, equals sign (=) indicates unchanged parameters.

### 4.1 Dural inflammation responding to the deficiency of the lymphatic system

Meninges represent the main site for the generation of migraine pain (Messlinger, 2009; Pietrobon and Moskowitz, 2013). This brain surrounding membranes comprise blood vessels, sensory nerve fibers and a high diversity of local immune cells (McIlvried et al., 2017; Olesen et al., 2009; Pietrobon and Moskowitz, 2013). Among the latter, lymphocytes and mast cells have been reported to be present in high amount dura mater membranes (McIlvried et al., 2017). In migraine-like conditions, meninges are involved in so-called ‘neurogenic inflammation’, which is induced and maintained by neuropeptides, such as CGRP, released from trigeminal nerve fibers (Eftekhari et al., 2013; Mason et al., 1984). CGRP is currently recognized as one of the central migraine mediator, with multiple targets including dural vessels (Goadsby et al., 1988), meningeal nerve fibers (Cady et al., 2011) and immune cells (Ottosson and Edvinsson, 1997). Interestingly, it was proposed, that impaired clearance of CGRP from perivascular space can be a reason for post-traumatic headache after TBI (Piantino et al., 2019). Surprisingly, here we demonstrate that the release of CGRP in the meninges from K14 mice lacking mLVs is lower compared to WT control group, suggesting that K14 animals are less prone to trigger migraine attacks via this signalling pathway.. Noteworthy, CGRP is vital for lymphatic capillary formation (in particular, alleviating the severity of postoperative lymphedema) (Matsui et al., 2019), suggesting a potential positive crosstalk between CGRP signalling and activity of the meningeal lymphatic system. As local trigeminal nerve fibers are the main source of meningeal CGRP (Eftekhari et al., 2013; Mason et al., 1984), we proposed that the lymphatic impairment causes either a reduced storage of CGRP in trigeminal nerve terminals or lowered meningeal innervation by ‘peptidergic’ nerve fibers (Keller and Marfurt, 1991).

Previous studies on patients have mainly focused on cytokine profiling in the systemic circulation (Fidan et al., 2006; Khaiboullina et al., 2017; Oliveira et al., 2017; Perini et al., 2005; Sarchielli et al., 2004; Yücel et al., 2016), whereas the production of cytokines in the meninges have not been fully characterized. Various immune cells can be identified in the meninges, including cells transported within the mLVs (McIlvried et al., 2017; Alitalo, 2011; Hayday et al., 2001). These cells constitute the primary and major source of multiple pro-inflammatory and anti-inflammatory cytokines. Here we demonstrate that in the meninges the levels of IL-6, IL-10, MCP-1, TNFα, IL-12p70, and IFNγ are detectable already at basal conditions. In particular, we found the presence of the pro-inflammatory cytokines TNFα and IL-6, known to be associated with migraine pathology (Bø et al., 2009; Khaiboullina et al., 2017; Perini et al., 2005; Sarchielli et al., 2006; Wang et al., 2015; Yücel et al., 2016). Notably, the level of IL-6 was exceptionally high but did not differ between the two genotypes. However, we found that the absence of functional mLVs was associated with the increased levels of the pro-inflammatory cytokine MCP-1 (Wood et al., 2014). MCP-1 is known as the activator of mast cells, provoking release of mast cells derived components (Lv et al., 2012). In addition, MCP-1 delays mast cell death (Lopes et al., 2019). This supports our finding of higher number of mast cells in meninges lacking mLVs. Notably, an increase in MCP-1 was reported in the cerebrospinal fluid of patients during a migraine attack (Bø et al., 2009). In contrast, the levels of the pro-inflammatory cytokine (IL-12p70) (Martins et al., 2011), which is typically released from macrophages and dendritic cells (Müller-Berghaus et al., 2004), were reduced in the meninges from K14 animals lacking mLVs. It has been shown that IL12-p70 plasma level was increased in migraine patients during the interictal phase (Oliveira et al., 2017) . The reduction of IL12-p70 in K14 animals lacking mLVs probably suggests a decreased inflammatory state in the meninges. Notably, mast cells can release IL12-p70 (Mukai et al., 2018) in response to CGRP-induced degranulation (Ottosson and Edvinsson, 1997; Theoharides et al., 2005). Our data showing decreased levels of CGRP in animals lacking mLVs may imply that the absence of a functional meningeal lymphatic system leads to a reduced tendency to develop a migraine through several downstream mechanisms controlled by this neuropeptide.

Extracellular ATP, acting primarily via P2X7 receptors, is a powerful trigger of inflammatory reactions involving different dural immune cells, including mast cells (Burnstock, 2016; Karmakar et al., 2016; Nurkhametova et al., 2019, 2018). Consistent with this view, we found that the stimulation of meninges with the P2X7 agonist BzATP stimulated the release of the pro-inflammatory TNFα, a promoter of trigeminal pain (Franceschini et al., 2013). Notably, ATP receptor stimulation resulted in the significant release of TNFα only in the meninges from WT animals, but not in the meninges lacking the mLVs. TNFα is the main migraine-related cytokine (Martami et al., 2018; Oliveira et al., 2017; Perini et al., 2005; Yücel et al., 2016). In line with this, TNFα has been reported as the agent, which can stimulate CGRP transcription (Bowen et al., 2006). Thus, the lack of increase in TNFα levels after ATP receptor stimulation is consistent with our other findings on the reduced level of CGRP in the K14 animals. Taken together, the lack of meningeal lymphatic system may induce a local shift in the balance of various humoral factors (i.e., CGRP and pro-inflammatory cytokines such as TNFα), suggesting a reduction in the neurogenic inflammation signalling.

### 4.2 Meningeal nociception and morphological properties of nerve terminals

ATP is a powerful trigger of meningeal nociception (Koroleva et al., 2019; Yegutkin et al., 2016). Notably, ATP can operate either directly via neuronal P2X3 receptors (Yegutkin et al., 2016), or by degranulating mast cells and release of serotonin, which acts via 5-HT3 receptors at meningeal nerve terminals (Koroleva et al., 2019). However, despite elevated mast cell levels found in K14 mice, these mechanisms do not lead to significant differences in spiking activity between the two genotypes.

Since CGRP is released by trigeminal afferents, we tested if the diminished CGRP release, observed in K14 meninges, resulted from the reduced density of trigeminal innervation of dura mater. To investigate this hypothesis, we analyzed several parameters of meningeal innervation. Meningeal sensory nerve fibers are characterized mainly by nociceptive Aδ and peptidergic C-fibers (Levy and Strassman, 2002), with the latter considered as the primary source of CGRP (Eftekhari et al., 2013). Analysis of total length of axonal fibers and of the number of terminal segments of meningeal afferents in the meninges of K14 mice did not show any difference compared to WT littermates. This suggests that the lower CGRP released in K14 mice is not affected by an altered density of meningeal innervation, but rather represents the different content of neuropeptides or a reduced functionality of the CGRP releasing machinery.

In conclusion, our data show that innate deficiency in the meningeal lymphatics (as observed in the K14 mice used in this study) results in the alteration of pro-inflammatory signals in the meninges, including the increased recruitment of mast cells and, on the other side, a decrease in the levels of the principal migraine-associated pro-inflammatory cytokines. This novel phenotype suggests the presence of modulatory (compensatory) mechanisms aimed at balancing the local inflammation in the meninges and limit the triggering of nociceptive signals.

### 4.3 Limitations of the model

To the best of our knowledge, this is the first study investigating the interactions between the meningeal lymphatic system, neurogenic inflammation and nociception, which can contribute to generation of migraine pain. However, the model used in our study has some limitations. First, all our experiments have been conducted in *ex vivo* conditions. In this preparation, the meningeal lymphatic system is not active, thus affecting the recruitment and trafficking of immune cells and clearance of migraine mediators. Second, our animals were not sensitized to mimic migraine attacks. Third, only ATP (although one of most powerful algogen) was tested as a trigger of the nociceptive response. Forth, it has been previously reported that K14 mice present alterations in both systemic (Thomas et al., 2012) and neuro-immunity (Wojciechowski et al., 2019), which involves mainly the T lymphocytes, which is only a subset of immune cells known to be involved in migraine (Nurkhametova et al., 2018).

Therefore, in future studies, our results needs to be confirmed in *in vivo* migraine models (e.g., nitroglycerin injection or CSD induction) (Demartini et al., 2019; Van Den Maagdenberg et al., 2004) to better understand the impact of the meningeal lymphatic system on migraine state.

### 4.4 Pathophysiological implications

Here we explored a possible contribution of mLVs in migraine pathology, given the generally accepted central role of the meningeal tissues in this disorder. Our main working hypothesis was that the lack of functional meningeal lymphatic system would affect the clearance of the dural environment, resulting in the exaggeration of neurogenic inflammation and nociception, and leading to more severe symptoms of migraine.

Indeed, we found that the lack of the meningeal lymphatic is related to pro-nociceptive changes in the trigeminovascular system relevant to migraine pathology (*i.e.*, enhanced expression of the pro-inflammatory cytokine MCP-1). However, we surprisingly also found that the expression of the key migraine mediator CGRP and of the pro-inflammatory cytokines TNFα and IL-12p70 was decreased. This suggest that the absence of mLVs caused not uniform, but rather more complex and eventually functionally opposite, changes in the inflammatory and nociceptive mechanisms in the meninges. Thus, the lack of mLVs, without specific promoters of activated genetic factors, is not *per se* the main cause of migraine state, likely due to the presence of the compensatory mechanisms primerely aimed at balance the local inflammation.

## 6 Author Contributions

NM, FMN and RashidG originally conceived the study; NM, FMN, RashidG elaborated the study design; NM, KK, RaisaG performed experiments; NM, AA, OG, AS, JT analyzed the data; NM, AA, OG, JT performed the statistical analysis; NM, FMN, AA, AS, FNM, RashidG drafted a significant portion of the manuscript and figures. TM supported the study and edited the manuscript. All authors contributed to manuscript revision, read, and approved the submitted version.

## 8 Acknowledgements

For help with cytokine release assay, we would like to thank Sara Wojciechowski, Dilyara Nurkhametova and Sanna Loppi. For meningeal immunostaining, we would like to thank Maria Vihma.

## 9 Data Availability

The raw data supporting the conclusions of this manuscript will be made available by the corresponding author, upon reasonable request, to any qualified researcher.

## Abbreviations

WT –: wild type mice
K14 –: K14-VEGFR3-Ig mice
mLVs –: meningeal lymphatic vessels

