## Supplementary for "The role of the meningeal lymphatic system in local inflammation and trigeminal nociception implicated in migraine pain"

### *Supplementary Material*

#### 1 Axon segmentation and innervation analysis

We developed a semi-automated segmentation technique to annotate axons and analyze the innervation in acquired 3D confocal microscopy images. To segment an axon, an expert inserts two points along the desired axon on the maximum projection of a 3D confocal image. Given the two points, a minimal path algorithm detected the centerline of that axon. We used active contours to segment the desired axon, initiating from the detected axonal centreline. In detail, we formed the 2D maximum intensity projection image along the direction of focal planes, i.e., $z$-axis, and enhanced axons using Frangi filter (Frangi et al., 1998). We normalized the enhanced images to the range [0, 1] and complemented the intensity of the enhanced images, such that axons appeared darker than their background, denoting the image as $P$. We used Benmansour and Cohen method (Benmansour and Cohen, 2011) to define the minimal path between two given points $p_1$ and $p_2$ as $C_{p_1,p_2}$, a path which minimizes an energy functional on $P$. The energy minimized by the minimal path is $E(\gamma) = \int_\gamma P(\gamma(s))\mathrm{d}s$, where $\gamma$ is a path parametrized along its length by $s$. To find $C_{p_1,p_2}$, we computed the minimal action map $U: \Omega \rightarrow \mathbb{R}^+$ associated to $p_1$ on the domain $\Omega \subset \mathbb{R}^2$ which is the minimal energy integrated along a path between $p_1$ and any point $x \in \Omega$. To compute $U$, we used the fast marching method (Sethian, 1996) with speed function $1 - P$. The map $U$ has only one local minimum, the source point $p_1$. Thus, the minimal path $C_{p_1,p_2}$ can be retrieved with a simple gradient descent on $U$ from $p_2$ to $p_1$, solving an ordinary differential equation with a standard numerical Runge-Kutta method. The minimal path $C_{p_1,p_2}$ is the axon centerline, because 1-$P$ was constructed using Frangi filter, admitting higher velocity in the center of the axon.

We used the extracted centerlines to segment the underlying axons using Chan-Vese active contours (Chan and Vese, 2001) (Matlab 2018b, Image Processing Toolbox, activecontour). For that, we initialized active contours from the extracted centerlines with speed function $1-P$. We set the contraction bias equal to -0.2, expanding the initial contours, and the maximum number of iterations to 10 to perform the curve evolution. The initial contours were deformed on the speed function to adapt to the shape of axons.

We also calculated the diameter of axons along their centerlines, as described in (Abdollahzadeh et al., 2019a). We calculated the length of the centerlines in 3D, by first projecting centerlines from $\mathbb{R}^2$ to $\mathbb{R}^3$, then smoothing the curves with an 11$^{th}$-order one-dimensional median filter and a shape-preserving piecewise cubic polynomial. We reported the sum of the length of all centerlines, excluding centerlines, which corresponded with axon bundles. The underlying segment of an axon bundle has a diameter thicker than a certain threshold, where we set the threshold to 8 µm. This threshold was acquired experimentally: the expert differentiated a few axonal bundles, where their average axonal diameter was measured 8 µm approximately. We quantified an innervation complexity value for each meningeal tissue by forming a graph from the union of the extracted centerlines as in (Abdollahzadeh et al., 2019b). We reported the number of graph terminal-vertices (endpoints) with the vertex degree equal to 1.

#### 2 K14-VEGFR3-Ig (K14) animals lacking mLVs

We used adult (30 – 40 g) male and female transgenic K14-VEGFR3-Ig (K14) mice lacking the meningeal lymphatic system (Aspelund et al., 2015; Mäkinen et al., 2001) and wild-type C57BL/6J littermates mice (WT). Figure 1 compares meninges from WT and K14 animals stained for LYVE-1 (red) and Prox-1 (green), confirming the lack of developed lymphatic vessels in the meninges of K14 animals.

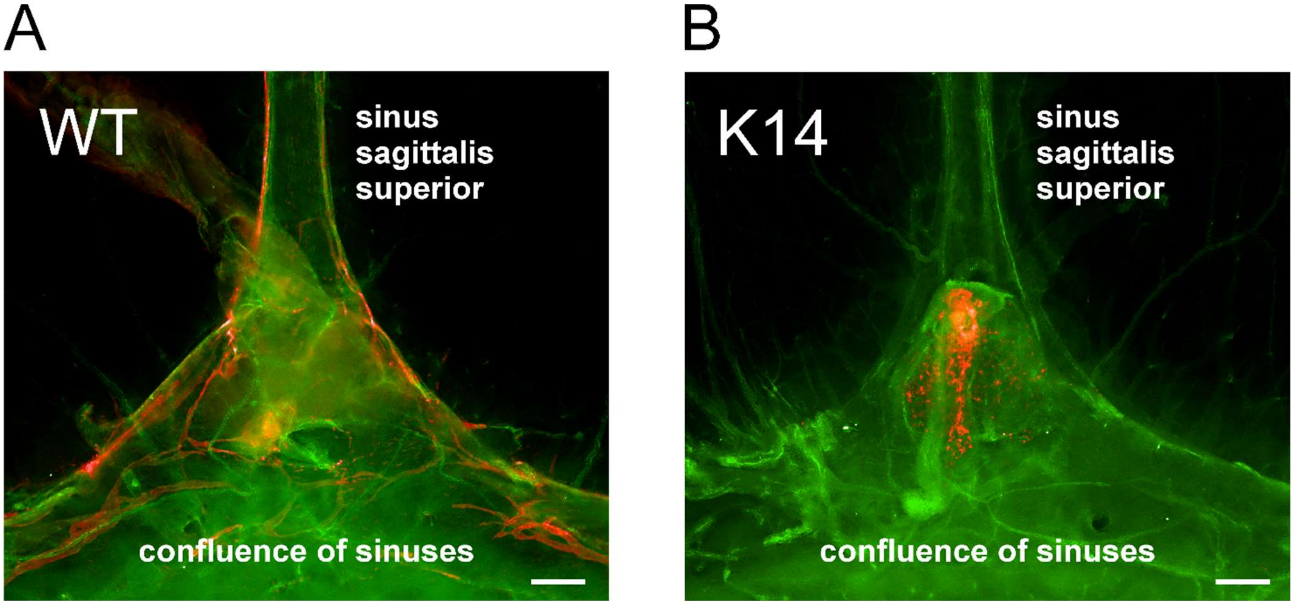


**Figure 1. Staining of meningeal lymphatic vessels in WT (A) and K14 (B) animals.** Panels present Prox-1 (green) and LYVE-1 (red) staining of lymphatic vessels. Scale bar – 200 μm.

Deeply anaesthetized mice were transcardially perfused with ice-cold 0.9% NaCl (6 ml ⁄ min, 6 min), followed by 4% paraformaldehyde (PFA) in 0.1 M sodium phosphate buffer, pH 7.4 (6 ml ⁄ min, 15 min). Skulls were post-fixed in 4 % PFA by immersion for 24 h at 4 °C. Meninges were then dissected from the skull and kept at 4 °C in PBS. The meninges were permeabilized with 0.3% Triton X-100 in PBS (PBS-TX) at room temperature (RT) and blocked with 5% donkey serum, 2% bovine serum albumin and 0.3% PBS-TX (DIM). The following primary antibodies were diluted in DIM, and incubated overnight at 4°C: rabbit anti–mouse PROX1 (1:200; #ab101851) and rat anti-mouse LYVE1 (1:300, MAB2125). After washing with PBS-TX, the meninges were incubated with fluorophore-conjugated secondary antibodies Alexa fluor594 Donkey anti-rat (1:500) and Alexa Fluor488 Donkey anti-rabbit (1:500) in PBS-TX overnight at 4°C, followed by washing in PBS-TX (5x) at RT. Fluorescent microscopy was acquired using an Axio Zoom.V16 fluorescence stereo zoom microscope (Carl Zeiss) equipped with ZEN 2012 software (Carl Zeiss) for images processing.
